# The chromatin remodelling factor Chd7 protects auditory neurons and sensory hair cells from stress-induced degeneration

**DOI:** 10.1101/2021.01.05.425431

**Authors:** Mohi Ahmed, Ruth Moon, Ravindra Singh Prajapati, Elysia James, M. Albert Basson, Andrea Streit

**Affiliations:** Centre for Craniofacial and Regenerative Biology, Floor 27 Tower Wing, Guy’s Hospital, King’s College London, London, SE1 9RT, UK; Wolfson Centre for Age-Related Diseases, Institute of Psychiatry, Psychology and Neuroscience, King’s College London, London, SE1 1UL, UK; MRC Centre for Neurodevelopmental Disorders, King’s College London, London SE1 1UL, UK

**Author notes:** Leukaemia and Stem Cell Biology Group, School of Cancer and Pharmaceutical Sciences, King’s College London, London, SE5 9NU, UK.

**Keywords:** CHARGE syndrome, Chromatin remodelling, Chromodomain helicase, Cochlea, Damage response, Degeneration, Epigenetics, Hair cells, Inner ear, Neurodevelopment, Neurodegeneration, Organ of Corti, Oxidative stress, RNA-binding proteins, Sensorineural hearing loss, Spiral ganglia neurons

## Abstract

Neurons and sensory cells are particularly vulnerable to oxidative stress due to their high oxygen demand during stimulus perception and transmission^1-4^. The mechanisms that protect them from stress-induced death and degeneration remain elusive. Here we show that embryonic deletion of the chromodomain helicase DNA-binding protein 7 (CHD7) in auditory neurons or hair cells leads to sensorineural hearing loss due to postnatal degeneration of both cell types. Mechanistically, we demonstrate that *CHD7* controls the expression of major stress pathway components. In its absence, hair cells are hypersensitive, dying rapidly after brief exposure to stress inducers, suggesting that sound at the onset of hearing triggers their degeneration. In humans, *CHD7* haploinsufficiency causes CHARGE syndrome, a disorder affecting multiple organs including the ear^5,6^. Our findings suggest that *CHD7* mutations cause developmentally silent phenotypes that predispose cells to postnatal degeneration due to a failure of protective mechanisms.

---

Sensorineural hearing loss (SNHL) is a common feature of CHARGE syndrome, affecting 50-70% of individuals^5,6^. Mice with heterozygous *Chd7* mutations are an excellent model for CHARGE and, like humans, exhibit SNHL^7-10^. *Chd7* plays an important role during neurogenesis both in the brain and the inner ear^9,11-16^. At embryonic day (E) 9.5-E10.5, neuronal progenitors are reduced in the inner ear of *Chd7*^*+/-*^ mutants, and Chd7 is necessary for their proliferation^9^. However, by E11.5, the number of neuronal progenitors is restored and inner ear neurons as well as the hair cells they innervate appear normal after birth^9,10^. The cellular function of *Chd7* and the mechanisms underlying SNHL have yet to be elucidated.

In the cochlea, inner hair cells are responsible for sound perception, while outer hair cells modulate the sound amplitude^17^. They are innervated by type I and type II spiral ganglion neurons, respectively, which project to the auditory nuclei in the brain stem^18^. In mice, hair cell specification mediated by the transcription factor Atoh1 occurs between E12.5 to E16.5^19,20^. However, their development continues postnatally^19^ and they reach maturity just before the onset of hearing between postnatal day 10 (P10) and P14^21^. To investigate its function in hair cells, we deleted *Chd7* using the hair cell-specific *Atoh1Cre* driver (*Atho1Cre/+;Chd7flox*). Surprisingly, loss of *Chd7* in newly formed hair cells does not affect their development: hair cells appear normal at P8 (n=10/10; Figure S1a, S1b). Thereafter, rapid degeneration of inner hair cells becomes evident between P10 (n=6/8) and P14 (n=15/16), while outer hair cells degenerate more slowly (Figures 1a-l, n, S1a, S1b). By P21, most inner hair cell nuclei are missing, pyknotic or fragmented, indicating progressive degeneration and cell death (n=8/8; Figures 1m, n, S1b). *Chd7* heterozygous mutants show equally severe phenotypes (Figure S1c) but at a reduced frequency (n=5/24). To establish when *Chd7* function is critical during hair cell formation, we used an inducible *Atoh1CreERT2*. Unlike *Chd7* deletion as soon as hair cells are specified, loss of *Chd7* after E16.5 does not cause postnatal hair cell degeneration (n=6/6; Figure S3b). These observations suggest that *Chd7* is required during hair cell development to maintain their survival upon the onset of hearing postnatally.

**Figure 1.**
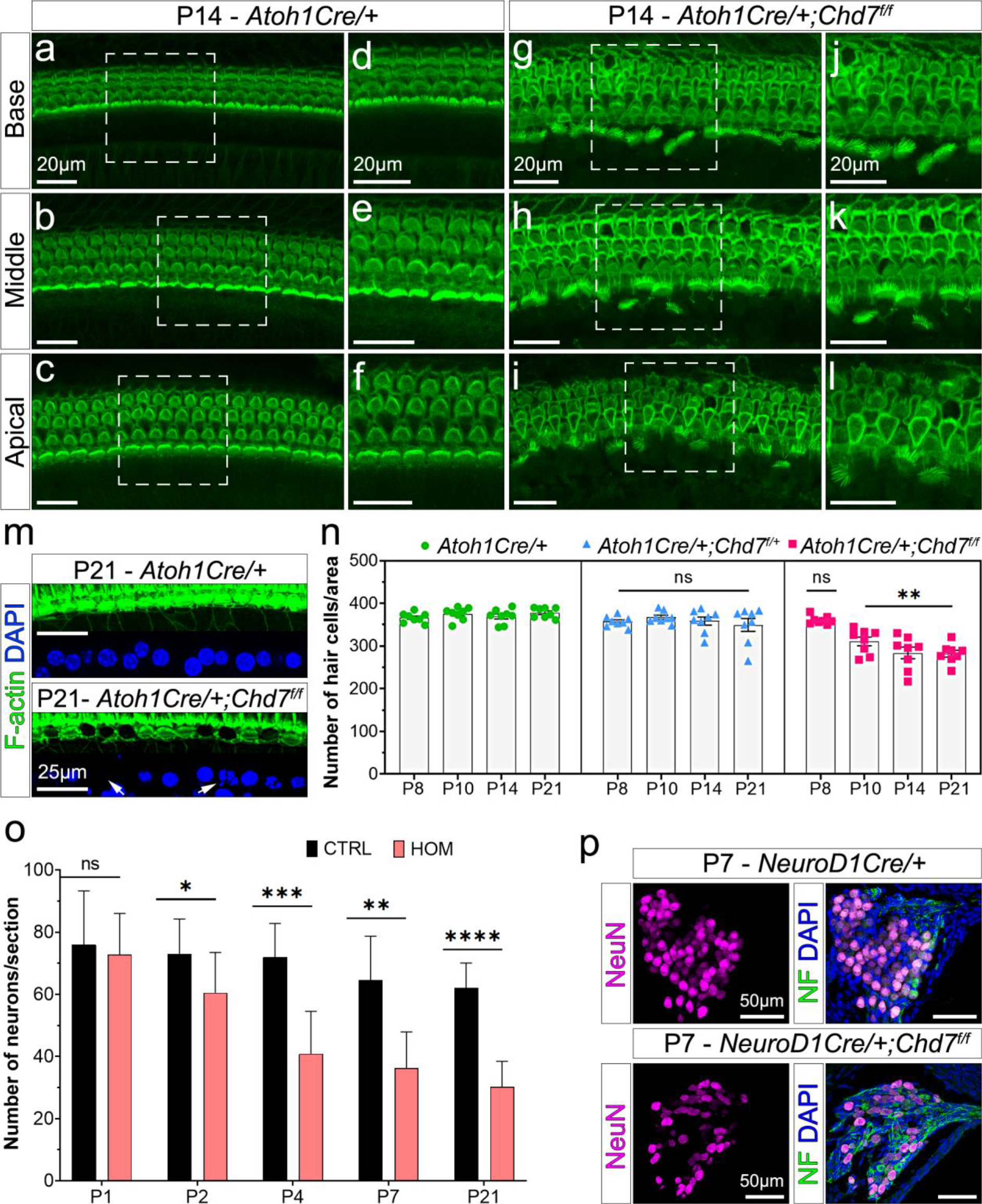
Postnatal degeneration of hair cells and neurons in *Chd7* mutants. **a-l**, F-actin-stained hair cells in the cochlea of control (a-f) and *Chd7* homozygous mutants (g-l) at P14. Dashed boxes in a-c and h-i indicate the zoomed regions shown in d-f and j-l. Scale bars = 20µm. **m**, Inner hair cells showing pyknotic, fragmented (arrow) and missing (arrow) nuclei in *Chd7* mutants at P21. Scale bars = 25µm. **n**, Average number of hair cells in three non-overlapping 200µm regions per base, middle and apical turn of each cochlea per animal (n=8 per genotype; each animal is represented by one circle, triangle or square). Separate inner and outer hair cell quantification is provided in Figure S2. Statistical significance was obtained by performing a nested one-way ANOVA and Dunnett’s multiple comparison test. ** P-value = 0.005. **o**, Average number of neurons in the spiral ganglion per section at different postnatal stages in control (CTRL) and *Chd7* homozygous mutants (HOM). **p**, Spiral ganglia neurons stained with NeuN and neurofilament (NF) at P7 in control and mutant animals (t test, P-values: *=<0.05, **<0.005, ***=<0.0005, ****=<0.00005). Scale bars = 50µm.

Postmitotic neural progenitors arise in the otic vesicle from ∼E9 onwards under the control of NeuroD1 and differentiate into spiral ganglion neurons by E14.5^22-25^. However, the peripheral auditory circuit is only established in the first 10 days after birth (P0-P10), prior to the onset of hearing^18,26,27^. To assess *Chd7* function in spiral ganglion neurons, we analysed *NeuroD1Cre/+;Chd7floxed* mutants. In homozygous mutants, ganglion size and neuronal numbers are indistinguishable from controls at P1 (n=3/3; Figures 1o, S2a), but neurons degenerate rapidly to 50% by P7 (n=3/3; Figure 1o, p). This phenotype is also observed in *Chd7* heterozygous mutants, although neurodegeneration occurs gradually with 50% loss by P21 (n=3/3; Figure S2b). Thus, *Chd7* controls the survival of a subset of spiral ganglion neurons. Like in hair cells, *Chd7* deletion at embryonic stages does not affect neuronal development but leads to delayed neurodegeneration postnatally.

To determine whether *Chd7* deletion results in hearing loss, we measured the auditory brainstem response (ABR) of 4- and 8-week-old mutant and control animals. Most *Atoh1Cre/+;Chd7flox* homozygous mutants exhibit severe-profound hearing loss across all frequencies (n=6/7; Figure 2a, S3a), while only 1/7 heterozygous mutants show a similar ABR profile (Figure S3a). In contrast, *NeuroD1Cre/+;Chd7flox* mutants exhibit moderate hearing loss (n=6; Figure 2b, S5a), presumably due to surviving neurons. Nonetheless, the ABR tests confirm that SNHL correlates with hair cell or neuronal degeneration.

**Figure 2.**
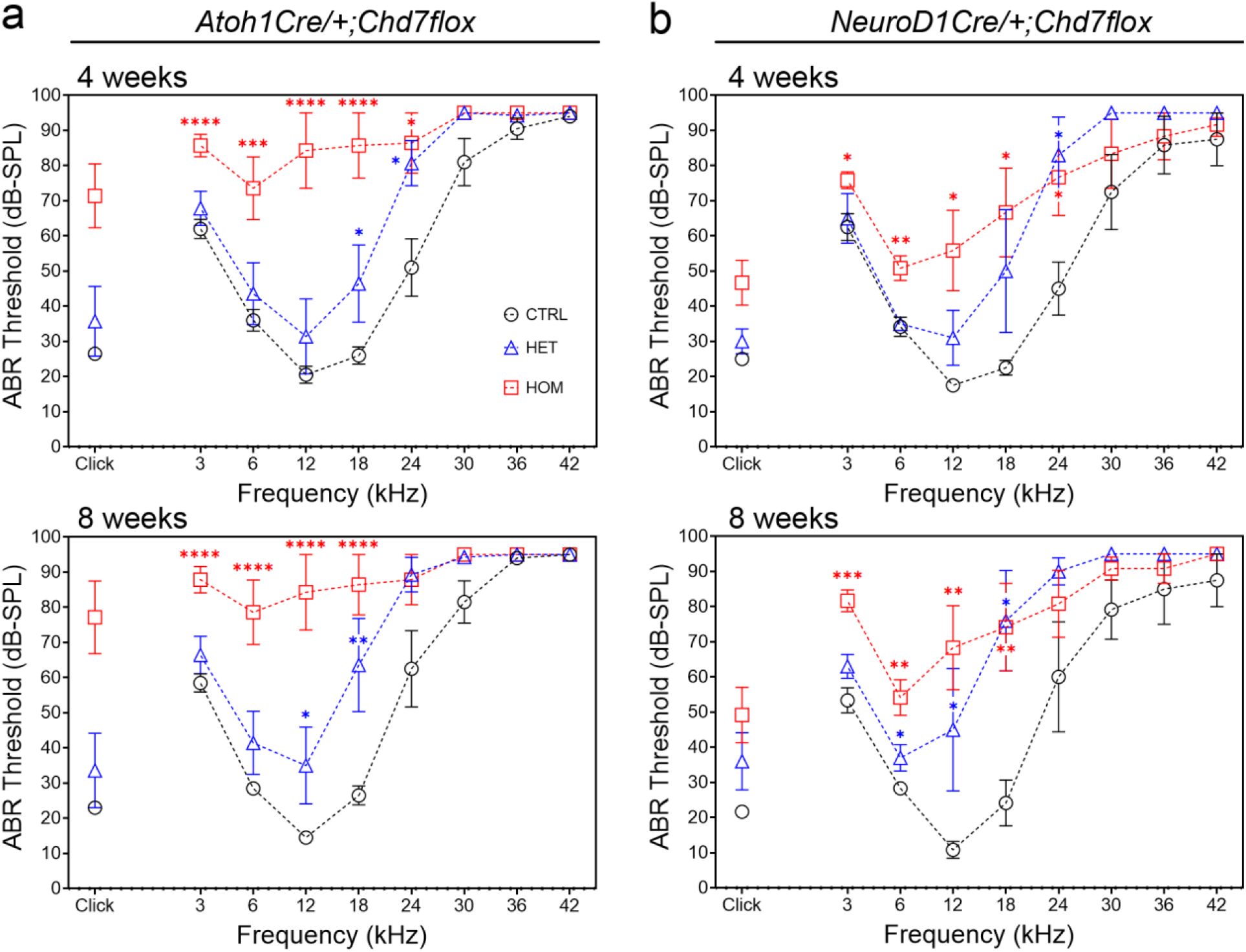
Chd7 mutants are hearing impaired. **a**, Auditory brainstem response (ABR) tests of *Atoh1Cre/+;Chd7flox* mutants and controls at 4 weeks and 8 weeks reveals profound hearing loss in homozygous and mild-moderate hearing loss in heterozygous *Chd7* mutants across all frequencies. **b**, ABR tests of *NeuroD1Cre/+;Chd7flox* mutants and controls at 4 weeks and 8 weeks reveals mild-moderate hearing loss in homozygous mutants. Frequencies where significant threshold elevations were observed are indicated by asterisks (P-values: *<0.05, **<0.005, ***<0.0005, ****<0.000005). See Figures S5 and S6 for ABR profiles of each mouse. Error bars represent the standard error of mean (see Figures S5 and S6). CTRL = control; HET = heterozygote; HOM = homozygote.

Chd7 controls transcription through regulation of chromatin architecture^13,15^, but how it exerts its function in auditory hair cells and neurons is poorly understood. We therefore examined the earliest transcriptional changes resulting from *Chd7* deletion by comparing gene expression of FAC-sorted hair cells or spiral ganglia neurons from mutant and control animals (Figure 3a, Table S1, S4). Differential gene expression analysis reveals significant changes (FDR ≤0.05, fold change >2) in 2437 transcripts in hair cells and 1653 transcripts in neurons (Figure 3b, Figure S4a-d, Tables S1-4). We validated expression changes of selected genes by qRT-PCR (Figure 3d) and protein by immunohistochemistry (Figure S5b). Analysis of Disease Ontology terms for all differentially expressed genes shows enrichment of *Chd7*-associated syndromes as well as hearing loss. Surprisingly, there is also a strong association with neurodegenerative diseases including dementia (Figure 3f, Tables S2, S4, S5, S6) pointing towards a common mechanism underlying these conditions.

**Figure 3.**
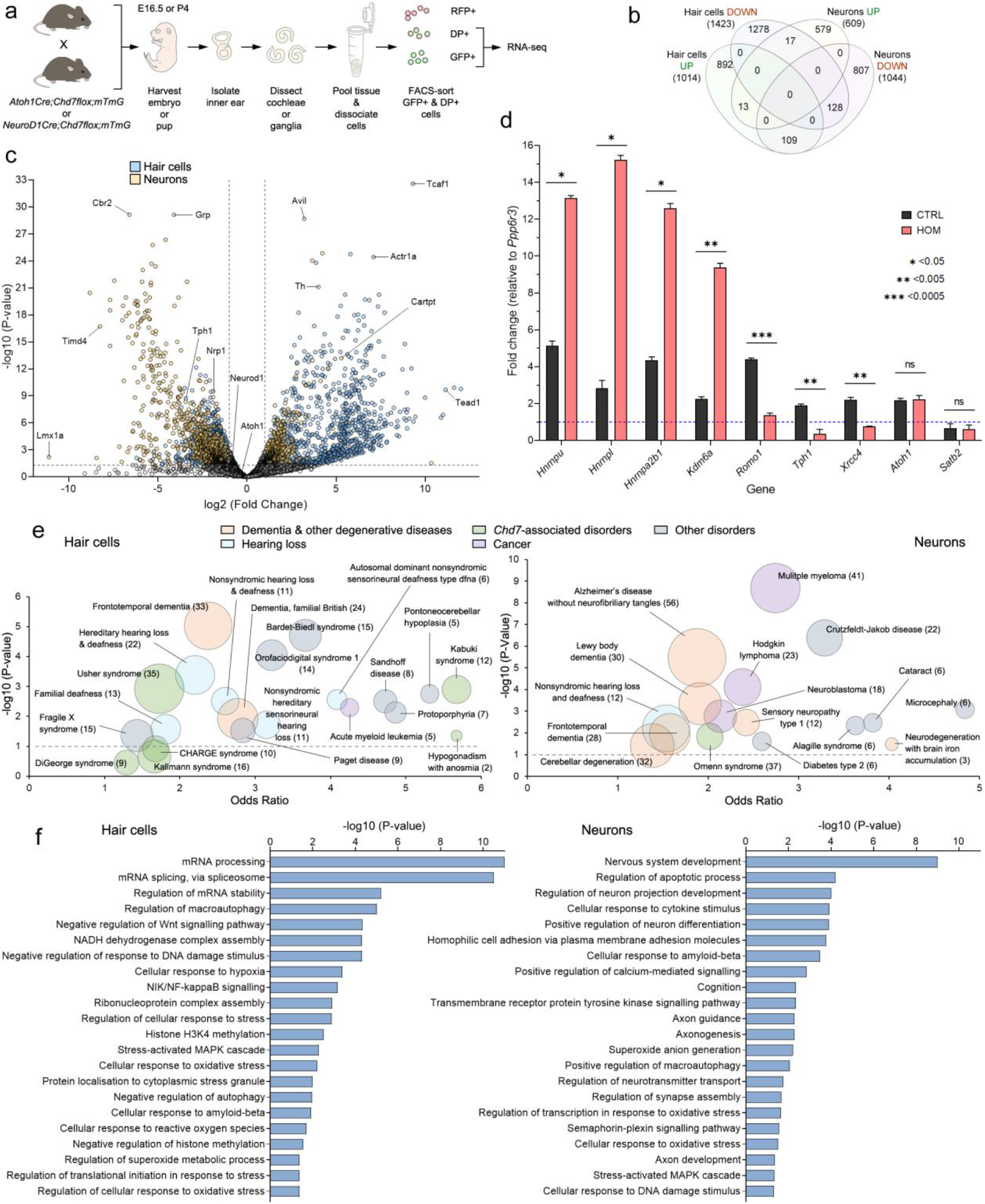
Transcriptome analysis of control and *Chd7* mutant hair cells and neurons reveals misregulation of cellular stress pathways. **a**, Schematic showing the experimental approach used for RNA sequencing. DP = double positive. n=6 cochleae or ganglia were pooled for RNA-seq in three independent experiments per genotype. **b**, Comparison of the number of differentially expressed genes between hair cells and neurons in homozygotes. **c**, Volcano plot displaying genes that are unaffected (grey) and significantly differentially expressed (adjusted P-value <0.05, fold change >2) in hair cells (blue) and neurons (amber). **d**, qPCR expression analysis of genes in controls and homozygous FAC-sorted hair cells at E16.5. Error bars represent the standard error. P-values: *=<0.05, **<0.005, ***<0.0005. ns = not significant. **e**, Plot of Odds ratio by -log10 of the P-value for human diseases identified by disease ontology. Complete disease ontology is provided in Tables S5 and S6. **f**, Gene ontology for differentially regulated genes. Complete gene ontology is provided in Tables S7 and S8.

Gene Ontology terms for RNA processing and stress pathways are strongly associated with all differentially expressed genes (Figure 3e, Table S3, S4), while RNA-binding proteins are among the most prominent transcripts deregulated by *Chd7* deletion (Figure 4). Indeed, in neuronal progenitors, Chd7 binds to many of their promoters (Figure S4e, Table S9; ref. 11) suggesting direct regulation. RNA-binding proteins are critical regulators of cellular stress, controlling the assembly and disassembly of stress granules^28-30^. As transient membrane-less compartments, they assemble in the cytoplasm under oxidative stress conditions to allow cells to survive, however their persistence triggers apoptosis^1,28-31^. The metabolic demands of sound detection and amplification elicits oxidative stress in neurons and hair cells that causes cell death unless tightly regulated^32,33^. We therefore tested the hypothesis that *Chd7* mutant hair cells are hypersensitive to oxidative stress by exploiting a cochlear explant system in which oxidative stress can be induced by treatment with aminoglycosides^4,34,35^. When P6 control explants from *Atoh1Cre/+;mTmG* mice are exposed to gentamicin for 5 hours (100µM), hair cells are intact along the entire length of the cochlea, as are untreated *Atoh1Cre/+;Chd7flox* homozygous mutant hair cells (n=5/5 each; Figure 5). In contrast, gentamicin treatment of mutant explants results in a reduction of hair cells by more than 50% across all regions of the cochlea (n=6/6; Figure 5). These findings show that *Chd7* mutant hair cells are hypersensitive to oxidative stress, causing degeneration in response to stress inducers. Thus, *in vivo* sound exposure at the onset of hearing may trigger cell death in Chd7-deficient hair cells. Our data suggest that SNHL in CHARGE syndrome may partly be due to mis-regulation of RNA-binding proteins as key regulators of stress granules thereby altering the response of neurons and hair cells to normal sound.

**Figure 4.**
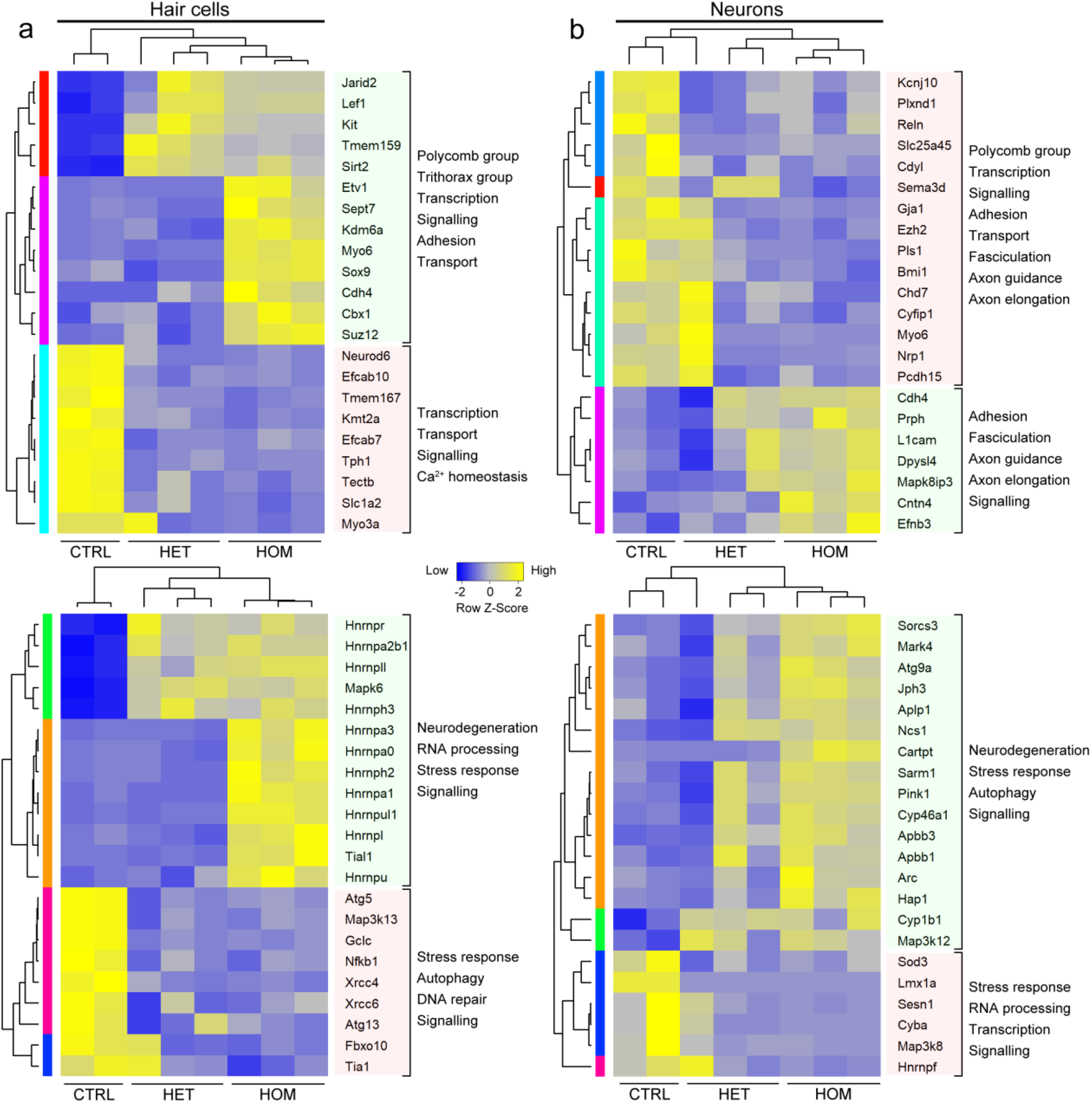
Chd7 regulates RNA-splicing and stress pathway genes. **a**, Heatmap of representative functional categories of differentially expressed genes in hair cells. **b**, Heatmap of representative functional categories of differentially expressed genes in neurons. Highlighted blocks of genes: green = upregulated; red = downregulated.

**Figure 5.**
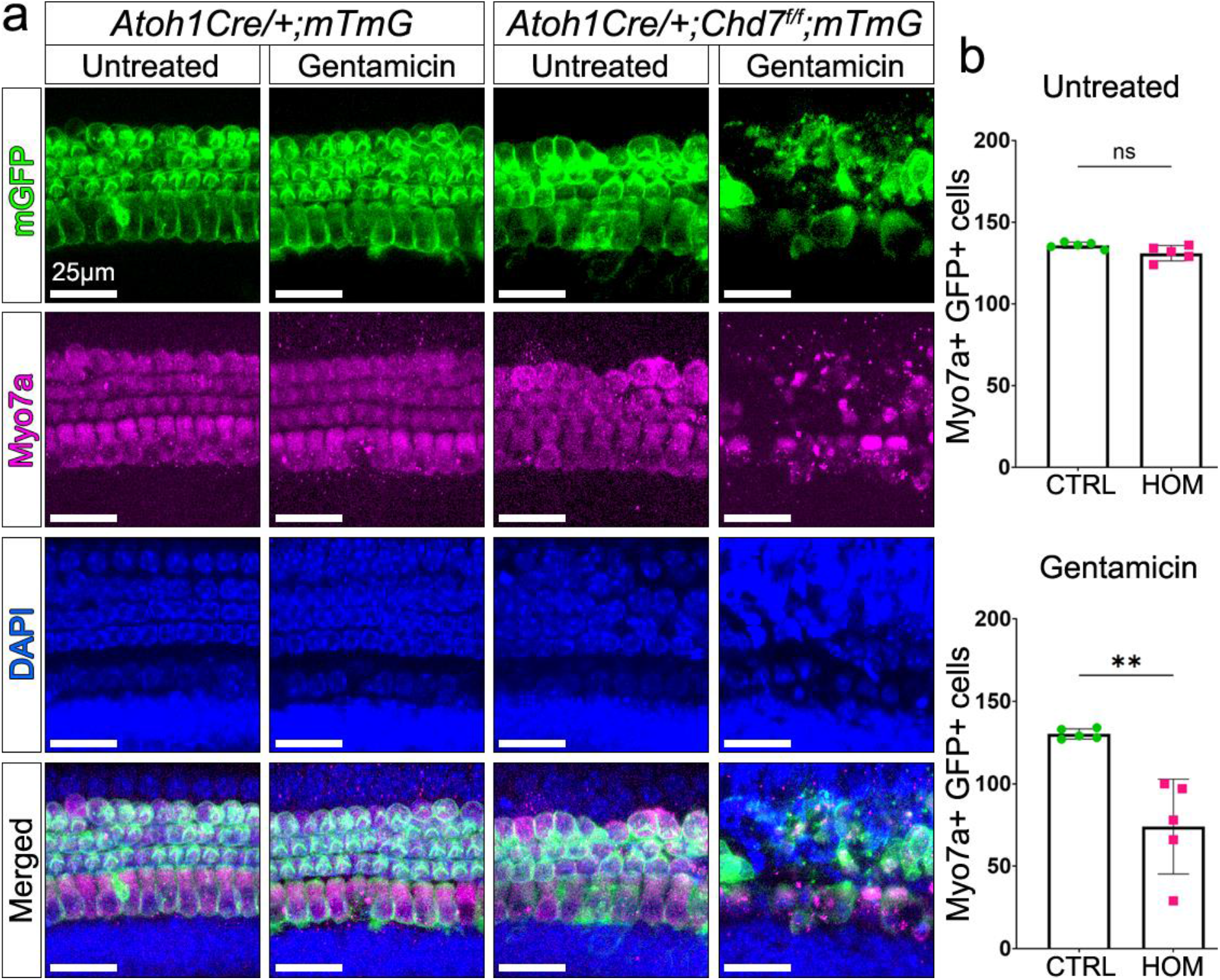
*Chd7* mutant hair cells are hypersensitive to stress. **a**, Cochlear explants of control and *Chd7* homozygous mutants were treated with gentamicin to induce oxidative stress. Rapid hair cell death is observed in mutants within 5 hours whereas control hair cells and untreated mutant hair cells survive. Green = Cre recombined cells expressing membrane GFP, magenta = all hair cells stained with Myo7a, blue = DAPI stained nuclei. **b**, Quantification of Myo7a+ and GFP+ hair cells per 200µm region in untreated and treated explants in both controls (CTRL) and mutants (HOM). Two-tailed unpaired t test shows significant difference between mutant and control. P-value = 0.0025. Scale bars = 25µm.

In summary, Chd7 emerges as a key coordinator of cellular stress proteins. Its embryonic deletion leads to an imbalance in stress pathways that does not affect normal development of neurons or hair cells. However, as cells mature and encounter environmental stress, they begin to degenerate. Our findings suggest that some neurodegenerative diseases arise from neurodevelopmental abnormalities that go undetected, and that SNHL may be an early indicator for these conditions.

## Acknowledgments

We thank Karen Steel and Claudio Stern for critical reading of the manuscript, Owen Harrison for excellent technical assistance, Zoe Mann and members of the Streit group for discussions. We thank Mary Beth Hatten for the *NeuroD1Cre* line. This work was supported by the MRC MR/R004625/1 and by Action on Hearing Loss (S39).

## Author contributions

MA conceptualised and designed the study together with AB and AS. MA performed most experiments and data analysis; RM analysed the neuronal *Chd7* phenotype, while EJ performed ABR tests. RP assisted in sequencing alignment and bioinformatics. MA and AS wrote the manuscript.

## Competing interests

The authors declare no competing interests.

## Methods

### Animals

The *Atoh1Cre (*B6.Cg-Tg(Atoh1-cre)1Bfri)^37^, *Chd7flox (*B6.tm1c(EUCOMM)Wtsi)^12^ and *NeuroD1Cre* (B6.Cg-Tg(NeuroD1-cre)RZ24Gsat^38^ mice were maintained on a C57BL/6J genetic background. The *mTmG (tm4(ACTB-tdTomato,-EGFP)Luo)*^39^ mice were maintained on a 129S6/SvEv background. The *Atoh1CreERT2* (Tg(Atoh1-cre/Esr1*)14Fsh, JAX #007684) used in Figure S4 was maintained on an FVB/NJ background. The *Chd7floxed* mice were crossed with the relevant Cre/reporter lines and backcrossed to C57BL/6J for three generations. All mice were maintained in either C57BL/6J or mixed genetic background. For tamoxifen-induced Cre recombination, a single dose of 20mg/ml tamoxifen (Sigma, T5648) dissolved in corn oil (Sigma, C8267) was administered to pregnant *Atoh1CreERT2*;*Chd7flox;mTmG* dams (80mg/kg of body weight) by intraperitoneal injection. To minimise abortion, 10mg/ml of progesterone (Sigma: P0130) was simultaneously administered at half tamoxifen dose (40mg/kg of body weight). One injection gave a consistent recombination efficiency of 95-99%. Upon Cre-mediated recombination, targeted cells expressed membrane GFP. All animal work was performed in accordance with UK Home Office regulations. Experiments were performed on male and female littermates and animals were randomly allocated to experimental groups.

### Immunohistochemistry

Dissected inner ear tissue was fixed in 4% paraformaldehyde (PFA) in phosphate buffered saline (PBS) and processed for whole mount immunostaining or frozen sectioning. For whole-mount immunostaining, following permeabilisation with 0.2% Triton X-100/PBS (3 x 10 minutes) and blocking with 0.2% Triton X-100/5% serum/PBS (1 hour), the cochleae were incubated overnight at 4°C in primary antibodies and then washed in 0.2% Triton X-100 (3 x 10 minutes). Fluorescent secondary antibodies were applied for 1 hour at room temperature. After staining with DAPI, the cochleae were washed extensively prior to mounting onto slides in 50% glycerol/PBS. For immunostaining on cryoprotected sections, following washes in PBS (2 x 10 minutes), permeabilisation in 0.1% TritonX-100/PBS (1 x 10 minutes) and blocking in 0.1% TritonX-100/5% serum/PBS (30 minutes), sections were incubated overnight at 4°C with primary antibodies. After several washes in 0.1% TritonX-100/PBS, the sections were incubated for 1 hour at room temperature with fluorescent secondary antibodies, subsequently washed in PBS, stained with DAPI and mounted onto slides with 50% glycerol/PBS. Primary antibodies used were: rabbit Myo7a (1:1000, Proteus, 25-6790); rabbit NeuN (1:1000, Abcam, ab177487); mouse NF-M (1:200, ThermoFisher Scientific, 13-0700); rabbit Sptbn1 (1:500, Bethyl Laboratories, A300-936A); rabbit Lmx1a (1:100, Abcam, ab139726); rabbit Epha3 (1:100, St John’s Laboratory, STJ110712). Secondary antibodies were: goat anti-rabbit Alexa Fluor 635 (1:500, Invitrogen, A31576); goat anti-mouse Alexa Fluor 488 (1:1000, Invitrogen, A11001). F-actin were stained with Phalloidin 488 (1:1000, Invitrogen, A12379) or 546 (1:500, Invitrogen, A22283).

### Auditory Brainstem Response (ABR)

ABR measurements were performed as described in ref. 40. An audiometric profile for each mouse at 4 and 8 weeks old was obtained across a range of sound frequencies (3, 6, 12, 18, 24, 30, 36 and 42 kHz). The mice were on a mixed genetic background (C57BL/6J x 129S6/SvEv). Statistical significance was obtained using Kruskall-Wallis non-parametric ANOVA and Bonferroni-corrected significance in GraphPad Prism 9.0.0.121.

### Isolation of hair cells and neurons by FAC-sorting

For RNA-sequencing, samples were collected for three biological replicates on independent occasions. E16.5 cochlear duct from *Atoh1Cre;Chd7flox* mice or P4 spiral ganglia neurons from *NeuroD1Cre;Chd7flox* mice were isolated from inner ears in cold L-15 medium (Thermofisher, 21083027). Tissues were cut into 3-6 pieces depending on stage and collected into low-binding tubes with L-15 on ice. Per experiment, a total of 6 cochleae or ganglia from three siblings were pooled into one tube. Excess L-15 was removed and 100µl of 20U/ml Papain (27mg/ml, Sigma, P3125) and 1U/ul RNase-free DNAse (Promega, M6101) in L-15 medium was added to each tube. Cells were dissociated at 37°C in a heated shaker, triturating using a filtered low-binding tip (Alpha Laboratories, LP200NFRS) every 5 minutes for a total of 40 minutes for hair cells and 1 hour for neurons. The dissociation reaction was stopped by adding 1:1 volume of prewarmed sample buffer (1% fetal bovine serum in L-15). Cells were strained using a 40µm nylon sterile cell strainer (Falcon, 352340) into a 50ml low-binding tube (VWR, 5250403) and transferred to a 5ml FACS tube (Falcon, 352235). DAPI (1mg/ml) was added (1:1000) immediately prior to FAC-sorting using the BD FACSAria sorters into 1.5ml low-binding tubes with 100µl of sample buffer. FAC-sorted cells were centrifuged at 4°C for 4 minutes at 8000 relative centrifugal force (Eppendorf centrifuge 54415R), frozen in liquid nitrogen and stored at −80°C or immediately processed for RNA extraction and first strand cDNA synthesis.

### RNA purification, library preparation and RNA Sequencing

FACS-sorted cells were processed using the NEB Monarch kit (T2010S/L) for polyA+ RNA isolation and NEBNext Single Cell/Low Input RNA Library Prep Kit for Illumina (E6420S/L) and NEBNext Multiplex Oligos for Illumina (Index Primers Set 1, E7335S/L) was used for library preparation (as per kit instructions). RNA and cDNA quality were analysed using Agilent Total RNA 6000 Pico or High Sensitivity DNA Assay on a Bioanalyser (Agilent, 2100). Additional library quality control was performed by the Oxford Genomic Centre at the Wellcome Centre for Human Genetics (funded by the Wellcome Trust, grant 203141/Z/16/Z) and sequenced using Illumina HiSeq 4000 75bp paired-end reads. Following quality control, paired reads were aligned to mouse MM10 genome assembly. Alignment was performed using HiSAT2 version 2.1.0 with the default parameters in Galaxy version 2.1.0^41,42^. To facilitate quantitative gene expression analysis, aligned reads for each sample were counted using featureCounts version1.6.4^43^.

### Differential gene expression analysis

Differential gene expression analysis was performed using DESeq2 version 2.11.40.6, applying parametric fit^44^. Prior to differential gene expression analysis, a number of filters were applied. We considered the RPKM values of genes that are not normally expressed in E16.5 hair cells (i.e., *Satb2*) or P4 spiral ganglia neurons (i.e., *Atoh1*) and removed all genes with an RPKM value equivalent to or less than *Satb2* or *Atoh1*. This resulted in 6910 transcripts for genes expressed in hair cells and 11293 genes expressed in neurons. We performed a pairwise comparison between controls and homozygotes and controls and heterozygotes. Considering an adjusted p-value (FDR) of ≤0.05 and linear fold change of >2 in either direction, we found a total of 2437 genes in hair cells (1014 upregulated, 1423 downregulated) and 1653 genes in neurons (609 upregulated, 1044 downregulated) that were differentially expressed in *Chd7* homozygous mutants compared to controls (Figure 3b, Tables S1-4, see Figure S7a-d for heterozygotes).

Gene Ontology and Disease Ontology analysis were performed both separately and together on up- or downregulated genes using the R interface (https://cran.r-project.org/web/packages/enrichR/vignettes/enrichR.html) to the Enrichr (https://maayanlab.cloud/Enrichr/) databases (see Tables S5-8 for specific databases for each analysis). Heatmaps and bubble plots were generated with the R packages *pheatmap* and *GOplot*. Volcano plots were generated in GraphPad Prism 9.0.0.121.

### Quantitative (q) RT-PCR

cDNA from RNA extracted from FAC-sorted hair cells or neurons were subjected to qPCR analysis with the AriaMx Real-Time PCR System (Agilent Technologies) using SYBR green and gene specific primers. Reactions were repeated in triplicates. Relative expression levels were calculated using 2^-ΔΔCT^ method using *Ppp6r3* as an endogenous housekeeping gene. Differences between experimental groups were compared using an unpaired two-tailed Student’s t-test and P-value ≤0.05 was considered statistically significant.

### Cochlear explants

Cochlear explants from postnatal day 6 *Atoh1Cre;mTmG* and *Atoh1Cre;Chd7f/f;mTmG* mice were cultured in MatTek dishes coated with CellTak (BD Biosciences). Culture medium comprised of L-15 medium (Thermofisher, 21083027), 5% FBS, 0.2% N2, and 0.001% ciprofloxacin. Explants were maintained at 37°C under 5% CO_2_ for 1 hr prior to gentamicin exposure. Explants were incubated with or without 100µM gentamicin for 5 hours. At 5 hours, explants were rinsed in PBS, fixed in 4% PFA (20 minutes at room temperature) and rinsed again in PBS (3 × 10 minutes) prior to immunohistochemistry. Experiments were performed on explants of basal and middle turns of the cochlea.

### Microscopy and Imaging

Confocal z stack images were obtained using a TCS SP5 confocal (Leica) microscope, projected using Fiji and further processed using Photoshop (Adobe). Figures were assembled in Photoshop.

### Quantification and statistical analysis

Statistical significance for hair cell and neuronal quantification was obtained by performing one-way ANOVA and Dunnett’s multiple comparison test or paired t-test. Differences between experimental groups in Figure 5 were compared using two-tailed unpaired t-tests. All statistical tests were conducted using Microsoft Excel and GraphPad Prism 9.0.0.121.

## Data availability

RNA-seq data are on the Gene Expression Omnibus, GEO: GSE163798.

## Notes

### Competing Interest Statement

The authors have declared no competing interest.

## References

1. Neal, M. & Richardson, J.R. Time to get personal: A framework for personalized targeting of oxidative stress in neurotoxicity and neurodegenerative disease. Curr Opin Toxicol. 7, 127–132 (2018).

2. Cheng, L. et al. Moderate noise induced cognition impairment of mice and its underlying mechanisms. Physiol. Behav. 104, 981–988 (2011).

3. Fujimoto, C & Yamasoba, T. Oxidative stresses and mitochondrial dysfunction in age-related hearing loss. Oxid Med Cell Longev. 2014, 582849 (2014).

4. Goncalves et al. Drug-induced stress granule formation protects sensory hair cells in mouse cochlear explants during ototoxicity. Sci Rep. 9,12501 (2019).

5. Vissers, L.E. et al. Mutations in a new member of the chromodomain gene family cause CHARGE syndrome. Nat. Genet. 36, 955–957 (2004).

6. Zentner, G.E., Layman, W.S., Martin, D.M. & Scacheri, P.C. Molecular and phenotypic aspects of CHD7 mutation in CHARGE syndrome. Am. J. Med. Genet. A. 152A, 674–686 (2010).

7. Bosman, E.A. et al. Multiple mutations in mouse Chd7 provide models for CHARGE syndrome. Hum Mol Genet. 14, 3463–3476 (2005).

8. Randall, V. et L. Great vessel development requires biallelic expression of Chd7 and Tbx1 in pharyngeal ectoderm in mice. J Clin Invest. 119, 3301–3310 (2009).

9. Hurd, E.A. et al. The ATP-dependent chromatin remodeling enzyme CHD7 regulates pro-neural gene expression and neurogenesis in the inner ear. Development. 137, 3139–3150 (2010).

10. Hurd, E.A. et al. Mature middle and inner ears express Chd7 and exhibit distinctive pathologies in a mouse model of CHARGE syndrome. Hear Res. 282, 184–195 (2011).

11. Engelen et al. Sox2 cooperates with Chd7 to regulate genes that are mutated in human syndromes. Nat. Genet. 43, 607–611 (2011).

12. Jones, K.M. et al. CHD7 Maintains Neural Stem Cell Quiescence and Prevents Premature Stem Cell Depletion in the Adult Hippocampus. Stem Cells. 33, 196–210 (2015).

13. Feng, W. et al. Chd7 is indispensable for mammalian brain development through activation of a neuronal differentiation programme. Nat Commun. 8, 14758 (2017).

14. Donovan, A.P.A. et al. Cerebellar Vermis and Midbrain Hypoplasia Upon Conditional Deletion of Chd7 from the Embryonic Mid-Hindbrain Region. Front Neuroanat. 11, 86 (2017).

15. Whittaker, D.E. et al. Distinct cerebellar foliation anomalies in a CHD7 haploinsufficient mouse model of CHARGE syndrome. Am J Med Genet C Semin Med Genet. 175C, 465–477 (2017).

16. Whittaker, D.E. et al. The chromatin remodeling factor CHD7 controls cerebellar development by regulating reelin expression. J Clin Invest. 127, 874–887 (2017).

17. Dallos, P. et al. Prestin-based outer hair cell motility is necessary for mammalian cochlear amplification. Neuron. 58, 333–339 (2008).

18. Koundakjian, E.J., Appler J.L., & Goodrich, L.V. Auditory neurons make stereotyped wiring decisions before maturation of their targets. J Neurosci. 27, 14078–14088 (2007).

19. Chen, P. et al. The role of Math1 in inner ear development: Uncoupling the establishment of the sensory primordium from hair cell fate determination. Development. 129, 2495–2505 (2002).

20. Ahmed, M. et al. Eya1-Six1 interaction is sufficient to induce hair cell fate in the cochlea by activating Atoh1 expression in cooperation with Sox2. Dev Cell. 22, 377–390 (2012).

21. Mikaelian, D., & Ruben R.J. Development of hearing in the normal Cba-J mouse: Correlation of physiological observations with behavioral responses and with cochlear anatomy. Acta Otolaryngol. 59:451–461 (1965).

22. Kim, W.Y. et al. NeuroD-null mice are deaf due to a severe loss of the inner ear sensory neurons during development. Development. 128, 417–426 (2001).

23. Evsen, L. et al. Progression of Neurogenesis in the Inner Ear Requires Inhibition of Sox2 Transcription by Neurogenin1 and Neurod1. J Neurosci. 33, 3879–3890 (2013).

24. Fritzsch, B., Pirvola, U. and Ylikoski, J. Making and breaking the innervation of the ear: neurotrophic support during ear development and its clinical implications. Cell Tissue Res. 295, 369–382 (1999).

25. Sanchez-Calderon, H., Milo, M., Leon, Y. and Varela-Nieto, I. A network of growth and transcription factors controls neuronal differentation and survival in the developing ear. Int. J. Dev. Biol. 51, 557–570 (2007).

26. Huang L.C. et al. Synaptic profiles during neurite extension, refinement and retraction in the developing cochlea. Neural Dev. B:38 (2012).

27. Michanski, S. et al. Mapping developmental maturation of inner hair cell ribbon synapses in the apical mouse cochlea. Proc Natl Acad Sci. 116, 6415–6424 (2019).

28. Gilks, N. et al. Stress granule assembly is mediated by prion-like aggregation of TIA-1. Mol Biol Cell. 15, 5383–5398 (2004).

29. Leeuw, F.D. et al. The cold-inducible RNA-binding protein migrates from the nucleus to cytoplasmic stress granules by a methylation-dependent mechanism and acts as a translational repressor. Aging Cell. 19, e13136 (2020).

30. Gopal, P.P. et al. Amyotrophic lateral sclerosis-linked mutations increase the viscosity of liquid-like TDP-43 RNP granules in neurons. Proc Natl Acad Sci. 114, 2466–2475 (2017).

31. Hansen, J.M., Jacob, B.R., & Piorczynski, T.D. Oxidative stress during development: Chemical-induced teratogenesis. Curr Opin Toxicol. 7, 110–115 (2018).

32. Ohlemiller, K.K., Wright, J.S., & Dugan, L.L. Early elevation of cochlear reactive oxygen species following noise exposure. Audiol Neurootol. 4, 229–36 (1999).

33. Henderson, D. et al. The role of oxidative stress in noise-induced hearing loss. Ear hearing. 27, 1–19 (2006).

34. Hirose, K., Hokenbery, D.M., & Rubel, E.W. Reactive oxygen species in chick hair cells after gentamicin exposure in vitro. Hear Res. 104, 1–14 (1997).

35. Towers, E.R. et al. Caprin-1 is a target of the deafness gene Pou4f3 and is recruited to stress granules in cochlear hair cells in response to ototoxic damage. J Cell Sci. 124, 1145– 1155 (2011).

36. Ohlemiller, K.K., & Gagnon M.P. Cellular Correlates of Progressive Hearing Loss in 129S6/SvEv Mice. J Comp Neurol. 469, 377–390 (2004).

37. Matei, V. et al. Smaller inner ear sensory epithelia in Neurog1 null mice are related to earlier hair cell terminal mitosis. Dev Dyn. 234, 633–650 (2005).

38. Gong, S., et al. Targeting CRE recombinase to specific neuron populations with Bacterial Artificial Chromosome constructs. Journal of Neuroscience 27, 9817–9823 (2007).

39. Muzumdar, M.D. et al. A global double-fluorescent Cre reporter mouse. Genesis. 45, 593–605 (2007).

40. Ingham, N.J., Pearson, S. & Steel, K.P. Using the Auditory Brainstem Response (ABR) to Determine Sensitivity of Hearing in Mutant Mice. Curr Protoc Mouse Biol. 1, 279–287 (2011).

41. Afgan, E. et al. The Galaxy platform for accessible, reproducible and collaborative biomedical analyses: 2018 update. Nucleic Acids Res. 46, W537–W544 (2018).

42. Kim, D., Langmead, B., & Salzberg, S.L. HISAT: a fast spliced aligner with low memory requirements. Nature Methods, 12, 357–360 (2015).

43. Liao, Y., Gordon, K.S, & Shi, W. featureCounts: an efficient general purpose program for assigning sequence reads to genomic features. Bioinformatics. 30, 923–930 (2014).

44. Love, M., Huber, W., & Anders, S. Moderated estimation of fold change and dispersion for RNA-seq data with DESeq2. Genome Biology. 15, 550 (2014).

