## Supplemental Figures for "The chromatin remodelling factor Chd7 protects auditory neurons and sensory hair cells from stress-induced degeneration"

### Supplementary Figures

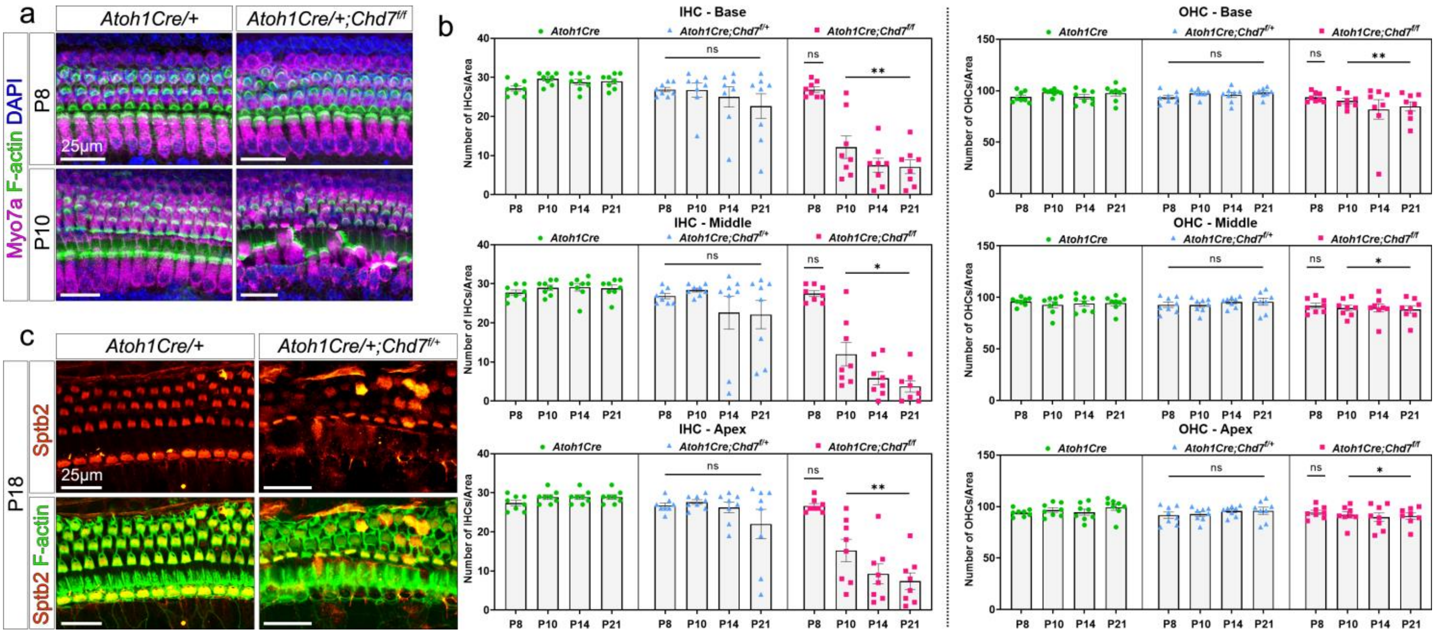


**Figure S1: *Atoh1Cre/+;Chd7flox* mutant hair cell phenotype. a**, Control images of *Atoh1Cre/+;Chd7f/f* mutants at P8 and P10. Myo7a labels all hair cells, F-actin labels the stereocilia on the apical surface of hair cells and DAPI stains the nuclei. **b**, Quantification of inner and outer hair cells in different regions of the cochlea. Nested one-way ANOVA and Dunnett's multiple comparison test was performed for statistical analysis. P-values: IHC base = 0.007; IHC middle = 0.01; IHC apex = 0.008; OHC base = 0.002; OHC middle = 0.04; OHC apex = 0.04. ns = not significant. **c**, P18 control and *Atoh1Cre/+;Chd7* heterozygous mutant stained with the cuticular plate and lateral wall maker β-II Spectrin (Sptb2) in combination with F-actin. Absence of Sptb2 in affected hair cells confirms the loss of the entire apical cap. Scale bars = 25µm.

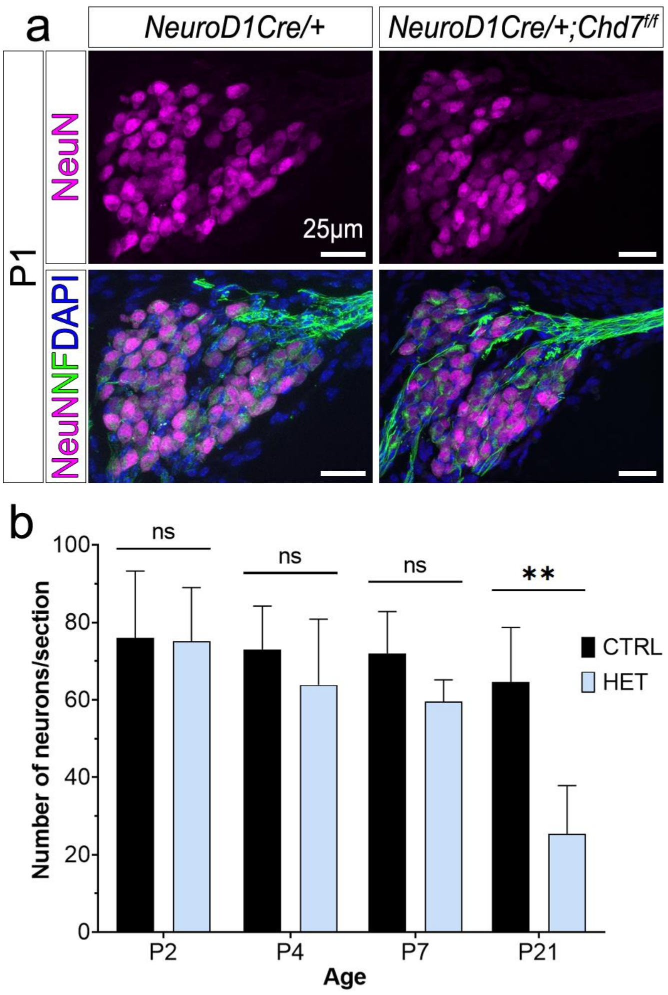


**Figure S2: *NeuroD1Cre/+;Chd7flox* mutant neuronal phenotype. a**, Control images of *NeuroD1Cre/+;Chd7f/f* mutants at P1. NeuN labels neuronal cell body and neurofilament (NF) labels axons. Scale bars = 25µm. **b**, Average number of neurons in the spiral ganglion per section at different postnatal stages in control and *NeuroD1Cre/+;Chd7* heterozygous mutants. ** P-value = <0.001.

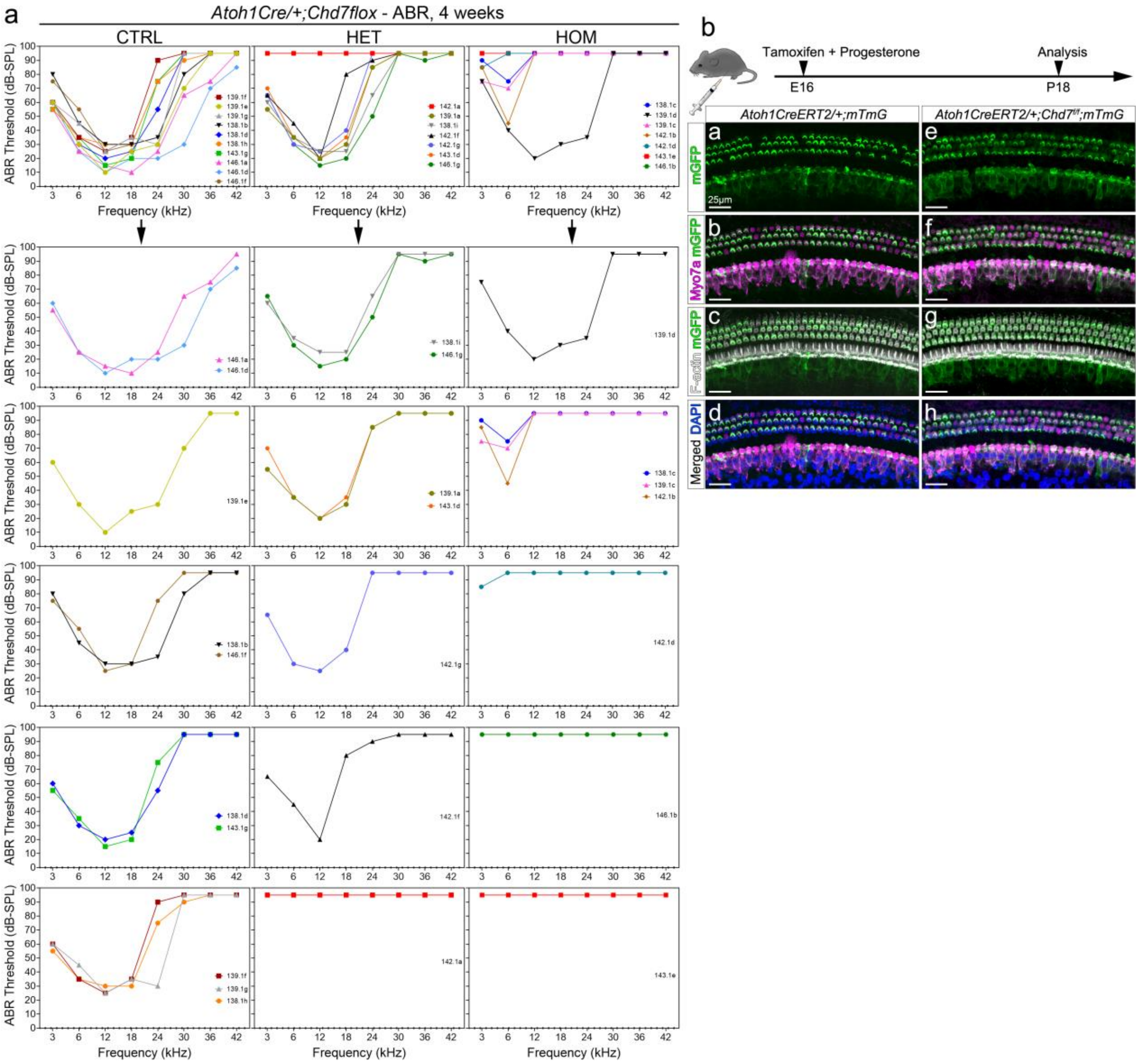


**Figure S3: *Chd7* deletion before E16 results in hearing loss but deletion after E16 does not cause a hair cell phenotype. a**, ABR thresholds of individual *Atoh1Cre/+;Chd7flox* animals*.* ABR thresholds for each mouse per genotype are represented by a coloured line (10 controls, 7 heterozygotes and 7 homozygotes). All mice were on a mixed (C57BL/6J x 129S6/SvEv) genetic background. 129S6/SvEv strain is expected to have accelerated age-related hearing loss (high frequency) at 1 month due to outer hair cell degeneration (ref. 36). Heterozygous mutants exhibit more variable ABR thresholds; however, 1 heterozygous mutant showed elevated thresholds across all frequencies identical to the most severely affected homozygous mutants. 6 homozygous mutants showed elevated thresholds across all frequencies with 1 mouse displaying an ABR profile similar to controls. The large standard deviation seen in Figure 2a is due to these outliers. **b**, Tamoxifen was administered to pregnant mice at E16 to induce Cre recombination (as indicated by mGFP). Cre recombination efficiency was ~95%. Analysis for hair cell phenotype was performed at P18. Note that there is no degeneration of hair cells.

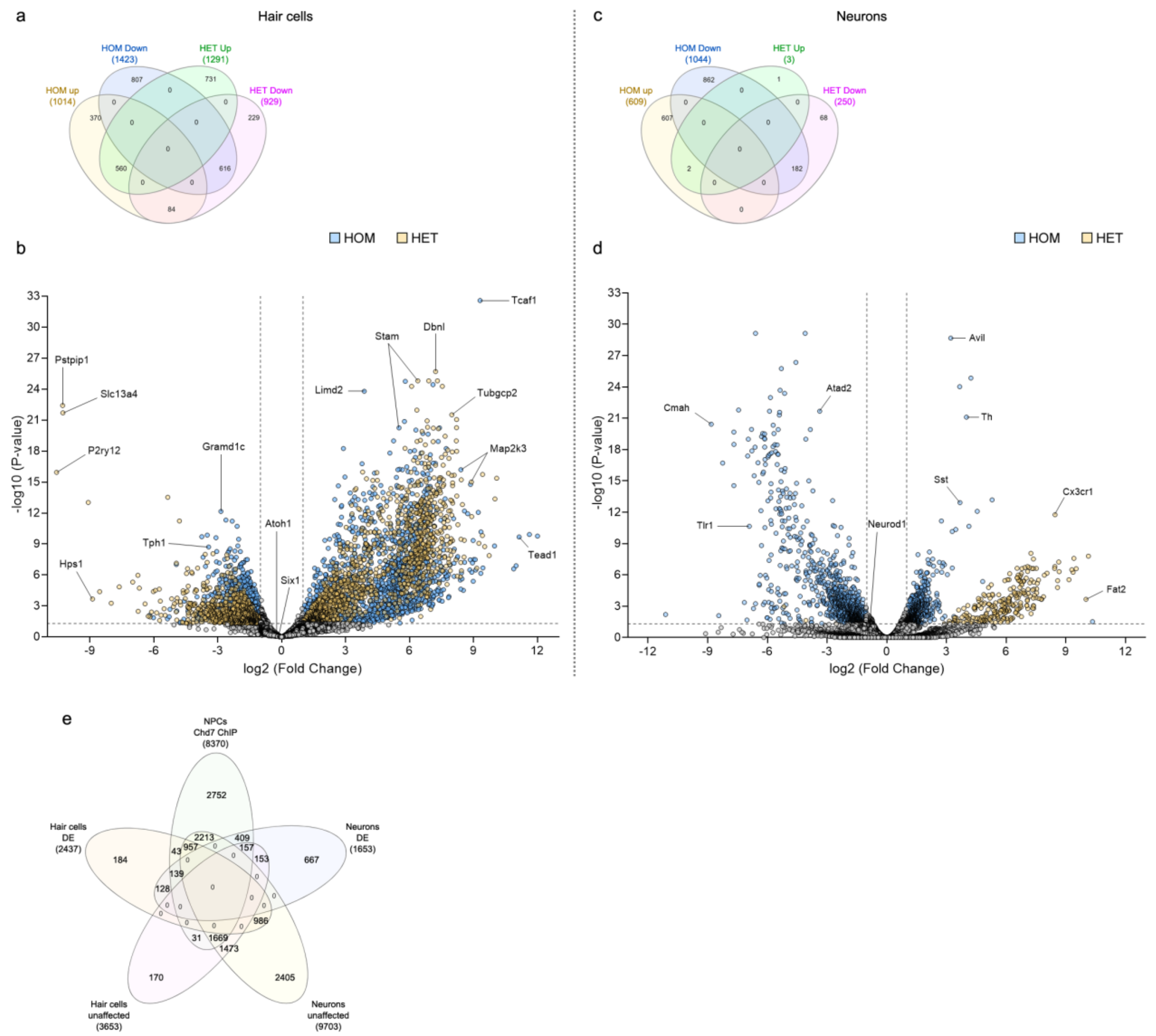


**Figure S4: RNA-seq analysis. a**, Comparison of the number of differentially expressed genes (DE) between homozygous and heterozygous mutants in hair cells. **b**, Volcano plot displaying genes that are unaffected (grey) and significantly differentially expressed (adjusted p-value <0.05, fold change >2) between *Atoh1Cre/+;Chd7flox* homozygous (blue) and heterozygous Chd7 mutants. **c**, Comparison of the number of differentially expressed genes between homozygous and heterozygous mutants in neurons. **d**, Volcano plot displaying genes that are unaffected and significantly differentially expressed (adjusted P-value <0.05, fold change >2) between *NeuroD1Cre/+;Chd7flox* homozygous (blue) and heterozygous Chd7 mutants. **e**, Chd7 ChIP-seq data from neural progenitor cells (NPCs; ref.

11) were compared with hair cells and spiral ganglia neurons RNA-seq data to identify genes directly bound by Chd7.

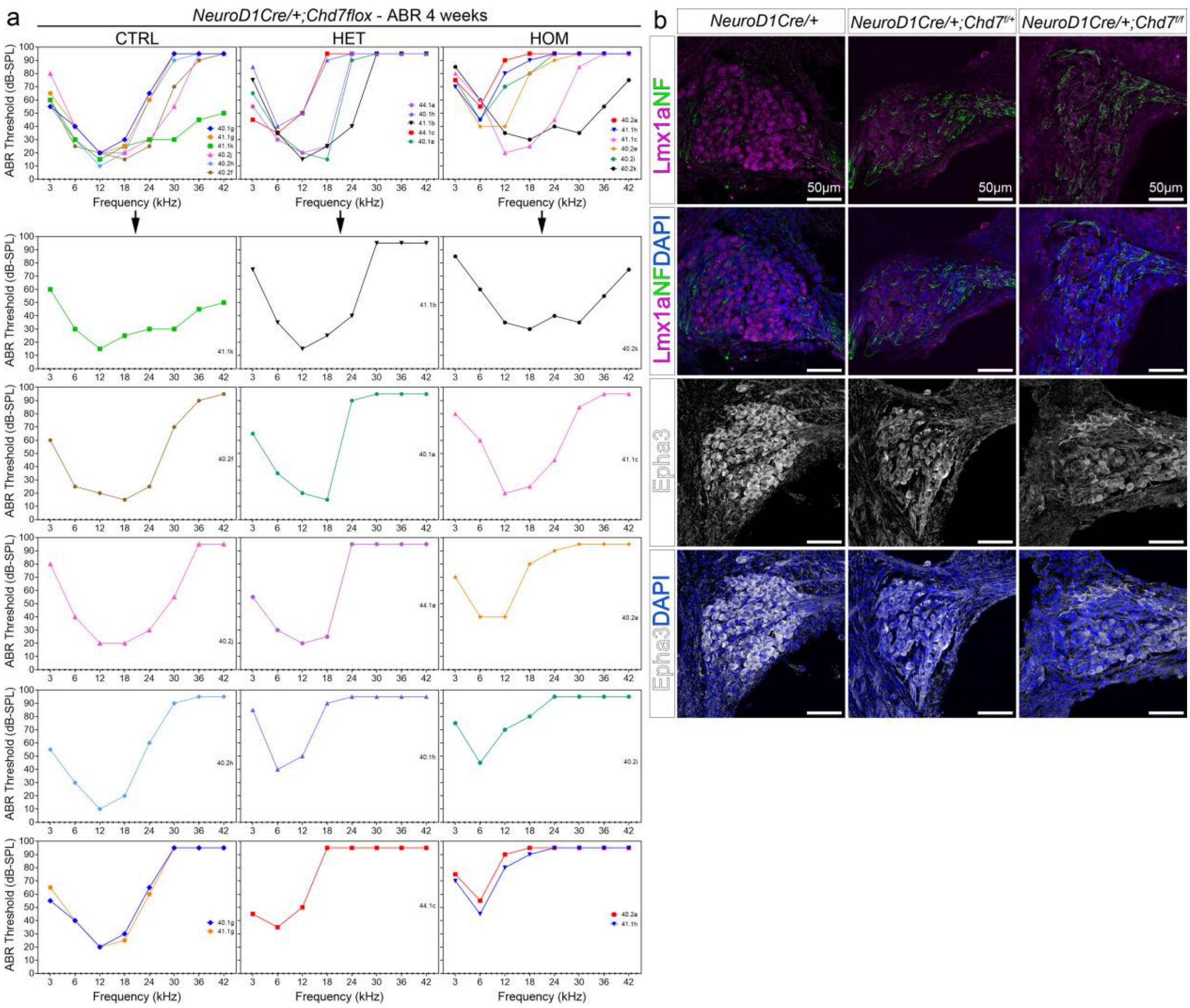


**Figure S5: ABR measurements and validation of RNA-seq data for selected proteins.**

**a**, ABR of individual *NeuroD1Cre/+;Chd7flox* animals*.* ABR thresholds for each mouse per genotype are represented by a coloured line (6 controls, 5 heterozygotes and 6 homozygotes). Both heterozygous and homozygous mutants show variable ABR thresholds but most mutants, particularly homozygotes, had elevated thresholds compared to controls. The large standard deviation seen in Figure 2a reflects this variability. **b**, Immunohistochemistry for Lmx1a (P7) and Epha3 (P4) confirms their downregulation in *NeuroD1Cre/+;Chd7flox* mutants. Markers and genotypes are indicated on each panel.

**Supplementary Tables**

Table S1. FAC-sorted hair cell RNA sample, library data and RPKM values

Table S2. Summary of differential gene expression (hair cells)

Table S3. FAC-sorted spiral ganglia neurons RNA sample, library data and RPKM values

Table S4. Summary of differential gene expression (neurons)

Table S5. Hair cells: Disease Ontology

Table S6. Neurons: Disease Ontology

Table S7. Hair cells: Gene Ontology

Table S8. Neurons: Gene Ontology

Table S9. Chd7 ChIP-seq (neural progenitors, ref. 11)
